# Relaxation of purifying selection suggests low effective population size in eusocial Hymenoptera and solitary pollinating bees

**DOI:** 10.1101/2020.04.14.038893

**Authors:** Arthur Weyna, Jonathan Romiguier

## Abstract

With one of the highest number of parasitic, eusocial and pollinator species among all insect orders, Hymenoptera features a great diversity of lifestyles. At the population genetic level, such life-history strategies are expected to decrease effective population size and the efficiency of purifying selection. In this study, we tested this hypothesis by estimating the relative rate of non-synonymous substitution in 169 species to investigate the variation in natural selection efficiency throughout the hymenopteran tree of life. We found no effect of parasitism, but we show that relaxed selection is associated with eusociality, suggesting that the division of reproductive labour decreases effective population size in ants, bees and wasps. Unexpectedly, the effect of eusociality is marginal compared to a striking and widespread relaxation of selection in both social and non social bees, which indicates that these keystone pollinator species generally feature low effective population sizes. This widespread pattern suggests specific constraints in pollinating bees potentially linked to limited resource availability and high parental investment. The particularly high load of deleterious mutations we report in the genome of these crucial ecosystem engineer species also raises new concerns about their ongoing population decline.

This article has been peer-reviewed and recommended by Peer Community in Evolutionary Biology https://doi.org/10.24072/pci.evolbiol.100120

## Introduction

The intensity of genetic drift experienced by a population depends on its effective population size, Ne (Wright 1931). Deleterious mutations reach fixation with a higher probability in small populations, that undergo more drift, than in large populations where purifying selection is more efficient. Ne is usually defined for any observed population as the theoretical census size an ideal Wright-Fisher population should have to show the level of drift displayed by the observed population (Wang et al. 2016). While different definitions of Ne exist depending on the field, it generally correlates negatively with any process breaking the assumption of panmixia, which underlies the Wright-Fisher model (i.e., population structure, homogamy, inbreeding…). Building on this knowledge, it has been proposed that basic traits influencing the reproductive output and mating choices of organisms, such as life-history traits, should correlate with their genome-wide deleterious substitutions rates. Several examples confirming these predictions have been uncovered in the last two decades: species generation time, longevity or body mass were found to be positively correlated with the genome-wide dN/dS (i.e., the ratio of the non-synonymous substitution rate to the synonymous substitution rate) (Nikolaev et al. 2007; Romiguier et al. 2014; Popadin et al. 2007; Figuet et al. 2016; Botero-Castro et al. 2017; Rolland et al. 2020). However, most known examples are clustered within a few vertebrate taxa: mainly mammals, birds or reptiles. To date, only few examples of such patterns have been found in invertebrates, which casts doubt on the existence of a general relationship between life history strategies and the efficiency of natural selection in Metazoa. Various reasons might explain the difficulty to demonstrate such relationships in invertebrates: There is relatively fewer genomic data available than in mammals or birds, and gathering life-history data in a large number of non-model invertebrates can be difficult as they have generally received less attention than vertebrates. Effective population size comparisons among invertebrate clades can also be particularly difficult, as the existence of reproductive systems such as haplo-diploidy, where the male is haploid, affect Ne estimations (Wang et al. 2016).

Among invertebrates, all Hymenoptera conveniently share the same haplo-diploid system, while displaying a particularly wide diversity of life-history strategies. Notably, they exhibit extreme lifestyles that can be predicted to strongly influence their reproductive output, and thus their long-term Ne. First, many species within this clade are parasites of plants (phytophageous) or parasitoids of other arthropods (Mayhew 2016), which could shape their demography, as population structure and size of the host can restrict that of the parasite or, especially, of the parasitoid (Mazé-Guilmo et al. 2016). Second, the hymenopteran order contains a large number of pollinators, such as bees, that are involved in keystone insect-plant mutualisms and strictly depend on a limited floral resource generally scattered in time and space. Because of the limited availability of resources required for their brood, pollinating bees have to invest a lot of time and energy in terms of foraging, nesting and food processing for their offsprings (Zayed et al. 2005; Zayed and Packer 2007), which should limit their reproductive output and population size. Supporting this hypothesis, high parental investment has been previously identified as a general proxy for low Ne in animals (Romiguier et al. 2014a). Third, eusociality, which is a rare lifestyle in animals, is relatively common in Hymenoptera with at least 9 independent appearances (Hughes et al. 2008). Because reproduction is typically monopolized by few long-lived reproductive individuals (Keller and Genoud 1997), a decrease in long-term Ne and the efficiency of natural selection is often thought to be a general consequence of eusociality (Bromham and Leys 2005; Romiguier et al. 2014; Settepani et al. 2016). Maintenance of high relatedness within low-Ne inbred groups has also been proposed as a prerequisite to the evolution of eusociality because it favors altruistic behaviors through kin-selection (Hamilton 1964; Husseneder et al. 1999; Hughes et al. 2008; Tabadkani et al. 2012). Ancestral population bottlenecks could thus be a typical feature of taxa in which eusociality frequently evolves, which is the case for several independent clades in Hymenoptera. Alternatively, it has been hypothesized that important population bottlenecks may be rare in Hymenoptera, because the associated loss of genetic diversity on single locus sex determination would lead to the costly production of sterile diploid males (Asplen et al. 2009; Rabeling and Kronauer 2013).

So far, few studies have investigated how Ne varies among Hymenoptera, and all were restricted to the effect of eusociality alone (Owen 1985; Berkelhamer 1983; Reeve et al. 1985; Bromham and Leys 2005; Romiguier et al. 2014; Imrit et al. 2020). Only recent studies with genome-wide datasets have detected associations between eusociality and decreases in Ne (Romiguier et al. 2014; Imrit et al. 2020), but these studies are typically restricted to few taxa compared to studies that rejected any significant effect (Bromham and Leys 2005). Disregarding the joint effect of other potential Ne determinants (e.g., body-size, parasitism, pollen-feeding, haplo-diploidy) may bias results and explain the discrepancy among studies with low vs high number of species comparisons.

Here, we tried to better assess the respective effects of potential Ne determinants in Hymenoptera. We used a phylogenomic dataset of 3256 genes in 169 species of Hymenoptera (Peters et al. 2017), including 10 eusocial species distributed among 4 independent origins of eusociality (Formicidae: 3 species; Polistinae/Vespinae wasps: 3 species; Stenogastrinae wasps: 1 species; Corbiculate bees: 3 species), 112 parasitic species and 32 solitary pollinating bees. We estimated mean genomic dN/dS for each species and compared these estimations between solitary and eusocial taxa, as well as between free and parasitic taxa (see figure S3),. We also correlated these to body size, life-history descriptor variables of parasitoids (see table S1 for details) and geographical range descriptors (see table S5 for details). We further confirmed that detected increases in dN/dS do correspond to relaxed purifying selection (and thus to drops in Ne) via specialized analyses that differentiate positive selection from relaxed purifying selection. Unexpectedly, we found that, instead of large species, parasites or eusocial taxa, pollinating bees display by far the lowest long-term Ne among Hymenoptera.

## Results

### dN/dS distribution across the Hymenoptera phylogeny

We estimated dN/dS in 3241 gene alignments of 169 species of Hymenoptera using the mapNH program (Romiguier et al. 2012; https://github.com/BioPP/testnh) from the testnh program suite (Dutheil and Boussau 2008; Guéguen and Duret 2017). We used the tree obtained by Peters et al. (2017) and its topology through all analyses to correct for phylogenetic inertia. As eusocial Hymenoptera are known to have high recombination rates (Wilfert et al. 2007; Sirviö et al. 2011; Wallberg et al. 2015; Jones et al. 2019), which in turn are known to inflate dN/dS when associated to biased gene conversion in vertebrates (Duret and Galtier 2009; Lartillot 2012; Galtier et al. 2018), we estimated dN/dS considering GC-conservative substitutions only. Estimated rates should, therefore, be impervious to the effects of biased gene conversion (Galtier et al. 2018). Average corrected genomic dN/dS values are displayed along the hymenopteran tree on figure 1 (see the distribution of uncorrected dN/dS values in figure S1). The largest and smallest mean ratios were inferred for Eucera nigrescens (0.1901) and Cimbex rubida (0.0684), respectively. As expected for conserved coding regions, the distribution of genomic dN/dS ratios is close to 0 (overall average of 0.0947±0.003sd), indicative of the large prevalence of purifying selection. We observed above average dN/dS ratios in 3 of the 4 available eusocial clades: Formicidae (0.1068 ± 0.0093sd, 3 species), Polistinae/Vespinae wasps (0.1033 ± 0.0088sd, 3 species), and the Apis/Bombus/Tetragonula clade (0.1086 ± 0.0352sd). This last clade of bees does not clearly stand out however, as most bees in the dataset (Anthophila, species characterized by pollen feeding of larvae: Apidae, Megachilidae, Halictidae, Colettidae, Andrenidae, and Melittidae) show high dN/dS ratios (0.1190 ± 0.0302sd, 41 species) with no dependence on their social organization. Finally, only two purely solitary taxa displayed comparable dN/dS ratios: Siricoidea (0.1025 ± 0.0251sd, 3 species) and Cynipoidea (0.1005 ± 0.0175sd, 5 species).

**Figure 1.**
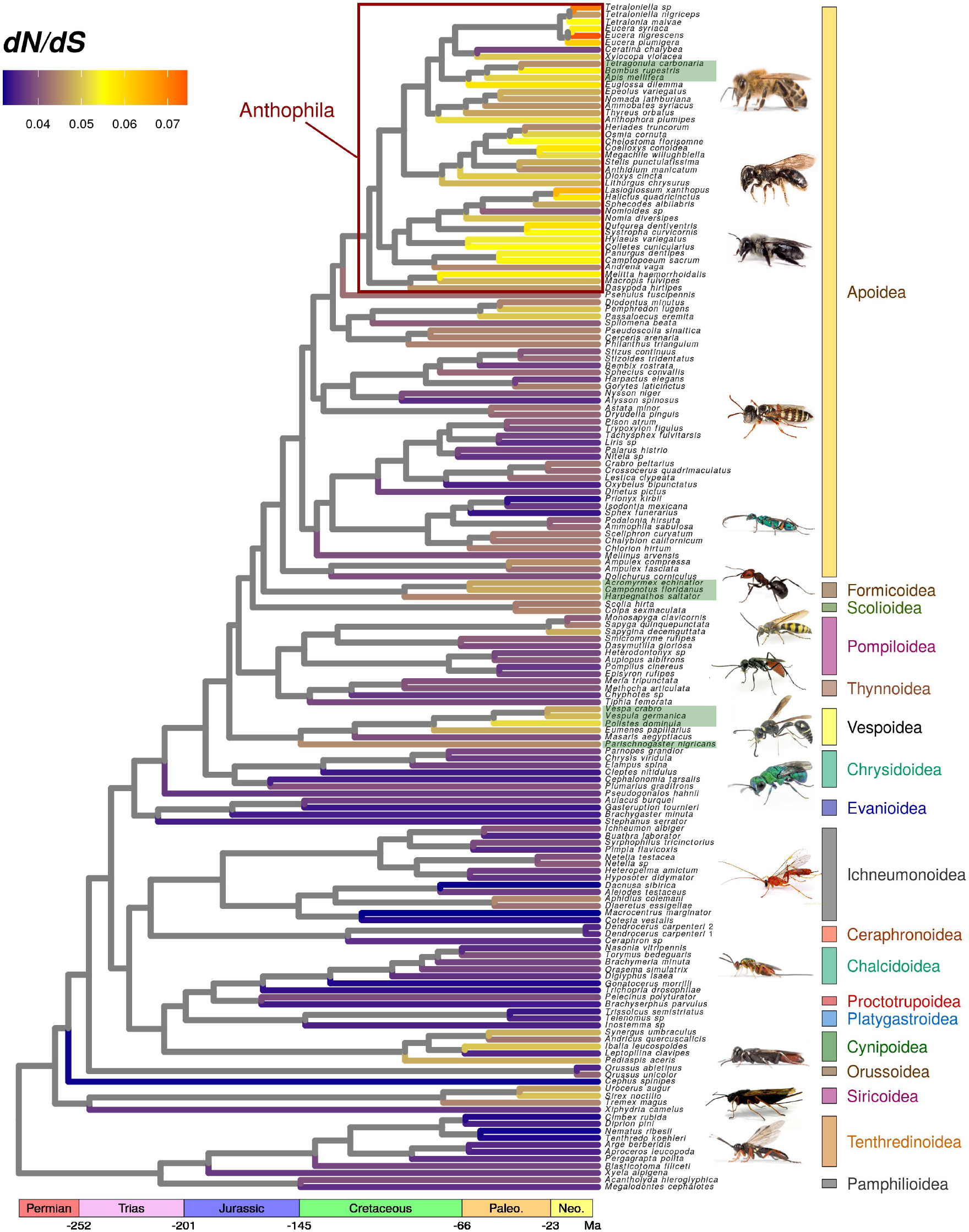
Corrected genomic dN/dS ratios for 169 species of Hymenoptera. dN/dS ratios estimated on terminal branches using 3241 genes are represented on the chronogram inferred by Peters et al. (2017). Green rectangles around labels indicate eusocial taxa.

We further used simple linear modeling to try and relate variation in dN/dS ratios to life history traits and geographical range descriptors. Phylogenetic independent contrasts were used to transform the data and account for phylogenetic relationships (Felsenstein 1985). We also used terminal branch length as a covariable in all models. This is because short terminal branches are known to bias dN/dS estimations upward as they yield more inaccurate estimations of this parameter, whose real value is often close to its zero boundary at a genomic scale. There is strong association between dN/dS ratios and branch length in this study (table 1). Variation in dN/dS estimation accuracy can also stem from variation in the number of genes available for each species. For example, four of the 10 available eusocial Hymenoptera (Apis mellifera and the three available ants), are species with published and annotated genomes (Consortium and The Honeybee Genome Sequencing Consortium 2006; Bonasio et al. 2010; Nygaard et al. 2011), and were used by Peters et al. (2017) as reference species for the identification of 1-1 orthologous genes, along with only one solitary reference species, Nasonia vitripennis (Werren et al. 2010). This translated into a relatively better power for gene prediction by Peters et al. (2017) in eusocial species, and thus into a significant (T=3.0567, df=9.3549, p-value=0.01305) over-representation of these eusocial species in alignments (mean number of alignments available per species: 2732.40 ± 88.09sd) compared to solitary species (2276.7 ± 90.74sd). To control for potential biases originating from varying precision in estimations, we replicated all the analyses of this study using a balanced subsampled dataset containing 134 alignments, each of them containing data for the same 88 species (most represented half of the species, referred later as the 88-species dataset). Unless specified otherwise, presented results were obtained with the full dataset. Average corrected genomic dN/dS estimated using the 88-species subsampled dataset are displayed along the Hymenoptera tree in figure S2.

**Table 1.** Linear modelling of corrected dN/dS ratios. Corrected dN/dS are obtained using 3241 genes and GC-conservative substitutions. Displayed results are obtained when simultaneously using all covariables inside a multiple linear model. Phylogenetic independent contrasts are used for all variables so as to account for phylogenetic autocorrelation.

| covariables | All samples<br>residual df = 127; $R^2=0.082$ | | | Non-Anthophila samples<br>residual df = 97; $R^2=0.117$ | | |
| --- | --- | --- | --- | --- | --- | --- |
| | $R^2$ | F | p-value | $R^2$ | F | p-value |
| branch length | 0.0347 | 4.8040 | **0.0302** | 0.0751 | 8.2422 | **0.0051** |
| adult size | 0.0085 | 1.1793 | 0.2795 | 0.0048 | 0.5363 | 0.4657 |
| Anthophila | 0.0381 | 5.2677 | **0.0233** | - | - | - |
| Eusociality | 0.0008 | 0.1232 | 0.7261 | 0.0372 | 4.0915 | **0.0458** |

### No effect of body size, parasitism and geographical range on relative protein evolution rates

Unlike findings in birds and mammals (Figuet et al. 2016; Botero-Castro et al. 2017), we found no significant effect of body size on dN/dS ratio in Hymenoptera (table 1). When testing for a difference in dN/dS ratios between parasitic (parasitoid or parasites) and free-living Hymenoptera (see table S3), we found a significant effect (df=167, F=46.327, p-val=1.715e-10, R2=0.217), but which completely disappeared when taking phylogeny into account (df=166, F=1.211, p-val=0.272, R2=0.007). We thus interpret this as being a confounding effect of sampling disequilibrium, as groups with elevated ratios completely lack parasites (with the exception of the cuckoo bumblebee Bombus rupestris and Sphecodes albilabris), and discarded this grouping from our models. We further tried to test for an association between dN/dS ratios of reproductive strategy and diet specialization within parasitoids using life-history and host range descriptors found in the literature (Traynor and Mayhew 2005a; Traynor and Mayhew 2005b; Jervis et al. 2003; Mayhew 2016), and summarized in table S1. However these descriptors were very seldom available for the species contained in the present phylogenomic dataset, forcing us to use genus-level averaging for both traits and dN/dS ratios. We detected no significant associations between average dN/dS ratios and life-history in parasitoids at the genus-level. We also tested for an association between dN/dS ratios and four proxies of species geographical range obtained using occurrence data available on the GBIF database. dN/dS ratios showed no significant correlation with mean latitude of occurrences, maximal distance between occurrences, or two additional estimators of species range (table S5).

### Anthophila bees and eusocial taxa reveal relaxed selection at the genomic scale

High dN/dS ratios in Anthophila bees is by far the strongest pattern observed in our results. Treating Anthophila as a covariable allows to significantly explain (df=167; F=175.84; p-val < 2.2.10-16) more than half the observed variation (R2=0.512). Despite Anthophila being only one monophyletic group, this effect is still present when accounting for phylogeny (table 1), and when accounting for sampling effort variation by using the 88-species subsampled dataset (table S3). This effect is strong enough to completely mask the effect of eusociality when using the full dataset. Indeed, the social status of a terminal branch significantly explains dN/dS variations in the dataset only if removing all Anthophila samples from the analysis. This is because eusocial corbiculate bees do not show any increase in dN/dS values when compared to other Anthophila. The increase of dN/dS in ants and eusocial wasps, remains significant when accounting for sampling effort variation by using the 88-species subsampled dataset (table S3).

To ensure that previous results stem from a relaxation of selection and not from strong positive selection, we applied the Hyphy RELAX procedure (Pond et al. 2005; Wertheim et al. 2014) on each available alignment separately. This procedure allows to formally test for selection relaxation by modelling the distribution of dN/dS ratios along the branches a phylogeny and by comparing the distribution fitted on a focal group of branches (eusocial taxa and Anthophila, alternatively) to the distribution fitted for the rest of the tree. Out of 3236 realized tests, 1743 (53.9%) detected relaxed selection on eusocial branches (including eusocial bees) and 184 (5.7%) detected intensified selection. Genes under relaxation of selection thus represent 90% of the genes for which a difference of selection efficiency between eusocial branches and focal branches could be detected. Results of a gene ontology enrichment analysis conducted with genes under intensified selection in eusocial species as focal genes are presented in table S4, but revealed no clear pattern. Using a conservative bonferroni correction for multiple testing in this procedure still leads to the detection of selection relaxation in 751 genes and of selection intensification in 28 genes. These results also hold if the more balanced 88-species subsampled dataset is used, as out of 134 alignments, 68 genes supported a relaxation of selection and 16 genes supported an intensification of selection. Moreover, the detected effect of eusociality does not seem to be driven by any over-representation of bees within eusocial species. The average number of eusocial bee sequences available for genes with relaxed selection (2.427 ± 0.018sd) is not different than within genes without relaxed selection (2.463 ± 0.024sd) (F=2.11; pval=0.146). These verifications are needed as bees experience an even stronger relaxation of selection. If this was apparent from simple modelling of genomic dN/dS ratios, it is made even more obvious by the application of the RELAX procedure with Anthophila branches as focal branches. Out of 3239 realized tests, 2000 (61.74%) detected relaxed selection on eusocial branches, while 294 detected an intensification of selection (9.07%). Using a conservative bonferroni correction for multiple testing in this procedure still leads to the detection of selection relaxation in 1210 genes and of selection intensification in 66 genes.

## Discussion

### Molecular consequences of body-size, parasitism and eusociality

Contrary to observed patterns in vertebrates (Nikolaev et al. 2007; Romiguier et al. 2014; Popadin et al. 2007; Figuet et al. 2016; Botero-Castro et al. 2017; Rolland et al. 2020), we did not detect any significant effect of body size on dN/dS in Hymenoptera. This suggests that the general association between body size and Ne observed in vertebrates is not universal in Metazoa, particularly in Hymenoptera. Parasitism is also not significantly associated to Ne decreases, which is surprising given the theoretical constrains imposed by the host (Papkou et al. 2016). This surprising negative result might be partly explained by the fact that Hymenoptera parasitoids lifecycle mostly requires a single insect host, a resource that might be not as limiting as parasites of vertebrates with complex lifecyles (Strobel et al. 2019).

We observed a significantly higher accumulation of non-synonymous substitutions in eusocial genomes, although the effect is relatively modest compared to the global pattern of increased dN/dS in pollinating bees (Anthophila). This increase can not be imputed to biased gene conversion, which is known to increase dN/dS by promoting the fixation of any G/C alleles (including deleterious alleles) (Rousselle et al. 2019), because our results are obtained using dN/dS ratios accounting only for GC-conservative substitutions. This result can not be imputed to positive selection either, as RELAX analyses detected relaxed selection on eusocial branches for more than half of the available alignments. This result supports the hypothesis of a relaxation of selection associated with eusociality through demographic effects. Long-lived reproductive female with delayed sexual maturity, as well as a biased sex-ratio and monopolization of the reproductive labor by few individuals, are typical features of eusocial species, which are bound to reduce effective population size. The hypothesis of a life-history effect matches well with the observation of a higher dN/dS in the highly eusocial formicoids ants Acromyrmex echinatior and Camponotus floridanus than in Harpegnatos saltator, which possesses a less complex social organization (Hölldobler and Wilson 1990).

These results are however to be taken with care, as the number of eusocial species in the dataset is low, and as no significant increase in dN/dS due to eusociality has been detected within bees. In this study, a choice was made not to increment the original dataset with additionnal eusocial species, because this addition would have introduced heterogeneity in sample treatment, and translated into new bias in the estimation of dN/dS ratios. It might be necessary to replicate our analyses using a separate, tailor-made and more exhaustive dataset in terms of eusocial species number in order to confirm the effect of eusociality on demography. Another exciting prospect will be to study Ne variation within eusocial groups. Ants, which display a variety of complexity levels in their social organisation, could represent an ideal model for a more quantitative approach (Bourke 1999), allowing to test for an effect of variation in eusocial characteristics of species on selection efficiency.

### Ecological and molecular predisposition to eusociality in bees

High genomic dN/dS ratios in all social and solitary bees unexpectedly appears as the major pattern of our results. Besides many independent transitions toward eusociality, Anthophila are characterized by their pollen-collecting behaviors which might explain our results. This dependence to large amounts of pollen to feed larvae is indeed believed to be a potential constraint on Ne, particularly in specialist species (Zayed et al. 2005; Zayed and Packer 2007). Pollen is a resource which is scattered in space and time and require a large energetic investment to come by and exploit (through progressive provisioning), thus constraining the very fecundity of females, which invest a lot of time and energy in their descent. Parental investment has already been highlighted as the major determinant of genetic diversity and long-term Ne in animal species (Romiguier et al. 2014). We suggest that high parental investment in pollinating bees might be a major factor limiting their Ne. This could in turn provide an explanation for the absence of differences between dN/dS ratios in social and solitary pollen-collecting species. Group-living might indeed represent a way to enhance the productivity of pollen collecting and metabolizing, thus compensating the decrease of Ne linked to eusociality in Anthophila.

Contrary to the general pattern in animals, pollinating bees appears as an exception and display higher species richness at high latitudes compared to tropics (Orr et al. 2020). This suggests that the diversification and origin of a pollinating bee lifestyle stems to environments with strong seasonality and important long-term climatic oscillations, which might have led to frequent bottlenecks in their population history. One previous study in Teleost fishes has shown that species of temperate regions display lower Ne than species of tropical regions (Rolland et al. 2020), while another found no such link across Metazoans (Romiguier et al. 2014b). Similarly to the latter, we found no associations between mean latitude (or other range descriptors) and dN/dS ratios in Anthophila (see table S5). However this result might be simply due to the massive over-representation of species from temperate regions in our dataset, and more thorough studies focusing on more tropical species will be necessary to draw any conclusions. In any case, specialized feeding on flowers appears here as a specialization to ecosystems with relatively low carrying capacity (Orr et al. 2020) requiring high parental investment for a scarce resource. Pollinating bees thus represent an ideal model to study the links between long-term demographics and seasonal variation in resource availability in temperate or arid environments. Anthophila could be also used to formally test whether the degree of specialization of a species towards one or a few plant species constrains Ne at the genomic scale in the general case (Zayed et al. 2005). Finally, bees might represent an opportunity to gain novel insights about the links between long-term demographics and characteristics more specific to Hymenoptera, such as nest parasite load (Wcislo 1987).

Interestingly, besides pollen-collecting, Anthophila (bees) is also the taxa with the highest number of independent origins of eusociality in the tree of life (Hughes et al. 2008). This suggests that low Ne is not only a consequence of group-living, but might also facilitate evolution toward eusociality. Supporting this hypothesis, low Ne due to intense inbreeding has been associated to communal behaviors in nesting and high parental care, two classical pre-requesites to an eusocial lifestyle (Hamilton 1964; Hussedener et al. 1999; Tabadkani et al. 2012; Wilson et al. 2008). Inbreeding also tends to increase within-group relatedness, which theoretically increases the benefit of kin selection, potentially favouring the emergence of eusociality (Hamilton 1964; Hussedener et al. 1999; Tabadkani et al. 2012; Kay et al. 2020; but see Nowak et al. 2010). Few genomic evidences supporting such a link have been observed so far. By showing a striking increase in dN/dS ratio in all Anthophila bees - the taxa concentrating more than half of the origins of eusociality in the tree of life - our results are the first genomic insight supporting the idea that low-Ne might have preceded and/or favoured evolution towards eusociality. As suggested previously in the literature, the evolution towards eusociality might have been favoured by the emergence of small groups of inbred individuals, despite the cost associated to genetic diversity loss at the sex determination single locus (Rabeling and Kronauer 2013).

Besides their implication regarding the evolution of eusociality, our results have important consequences for the conservation field. Pollination has been found to rely heavily on wild and domesticated bees, which ensure the majority of animal-mediated pollination of wild and domesticated plants in most ecosystems (Winfree 2010). Our finding of particularly high deleterious substitution rates within this group raises the additional concern that bee species might be especially sensitive to any further population decline, which are already known as particularly alarming (Murray et al. 2009; Arbetman et al. 2017; Powney et al. 2019).

## Conclusion

This study brings supplemental genomic evidence supporting an association between eusociality and reduced effective population size. We thus bring further support to the hypothesis that the extreme life-history traits of eusocial species constrain their molecular evolution. More interestingly, the surprisingly massive and widespread reduction of selection efficiency in both eusocial and solitary bees suggests unexpectedly high constraints of a pollinator lifestyle, potentially linked to limiting resource and high parental investment. This also brings genomic support to the hypothesis that some ecological characteristics associated with low Ne might have facilitated evolution towards eusociality. Altogether, this study suggests that, conversely to vertebrates, purifying selection efficiency in invertebrates is more constrained by lifestyle and ecology than simple body size.

## Data accessibility

The original dataset of Peters et al. (2017), with alignments and trees, is provided by the its authors at http://dx.doi.org/10.17632/trbj94zm2n.2. Detailed tables containing data used for this paper as well as obtained results are available at Zenodo.org : https://zenodo.org/record/3999857#.X0UsBBk6-it.

## Acknowledgements

We thanks Nicolas Galtier for advices during the writing of the manuscript and Laurent Keller for useful discussions. Version 5 of this preprint has been peer-reviewed and recommended by Peer Community In Evolutionary Biology (https://doi.org/10.24072/pci.evolbiol.100120)

## Materials and Methods

### Genetic data

Data was downloaded from the authors’ online repository (http://dx.doi.org/10.17632/trbj94zm2n.2). It originally contained nucleotide and amino-acids multi-sample alignments for 3256 protein coding genes predicted to be 1-1 orthologs in 174 species (see Peters et al. 2017 for details about the production of these alignments), 5 of which are outgroups to the Hymenoptera (2 Coleoptera, 1 Megaloptera, 1 Neuroptera and 1 Raphidioptera), and 10 of which are eusocial species. The latter belong to 5 independent eusocial clades: corbiculate bees (Tetragonula carbonaria, Bombus rupestris and Apis mellifera), ants (Acromyrmex echinatior, Camponotus floridanus and Harpegnathos saltator), Polistinae/Vespinae wasps (Vespa crabro, Vespula germanica and Polistes dominula), Stenogastrinae wasps (Parischnogaster nigricans).(Cardinal and Danforth 2011). The data also contained the trees inferred using this data by the original authors. We used the dated chronogram inferred by the authors using amino-acid data throughout this study. This tree corresponds to their main results and is contained in the file dated_tree_aa_inde_2_used_in_Fig1.tre available on the authors’ online repository.

### Data cleaning

Each amino-acid alignment was first checked for potential false homology using HmmCleaner (Di Franco et al. 2019; Philippe et al. 2017) with default settings. The resulting maskings were then reported on corresponding nucleotide sequences using the reportMaskAA2NT program from the MASCE program suite (Ranwez et al. 2011). At this point, we discarded individual sequences containing less than 50% of informative site within one alignment.

### dN/dS ratios estimation

Cleaned alignments were then used, along with the tree topology inferred by Peters et al. (2017) and the mapNH binary (Romiguier et al 2012; https://github.com/BioPP/testnh), to estimate synonymous and non-synonymous substitution rates along the branches of the Hymenoptera tree. MapNH allows a fast estimation of those rates by using tree-wide parameters obtained a priori by fitting a homogeneous model (YN98) to the data with the help of bppml (Dutheil and Boussau 2008), to parsimoniously map observed substitutions to the supplied topology. Estimated substitution counts for specific branches, obtained separately for each alignments, can then be summed to obtain genome-wide substitution rates. We used this method to obtain dN/dS ratios of terminal branches, susceptible to carry information about the long-term drift regime of extant lineages. 15 alignments did not contain enough data to allow correct convergence of the homogeneous model needed by mapNH.

### Controlling for biased gene conversion

We produced a corrected dN/dS using only GC conservative substitutions to estimate dN/dS. This was achieved using a custom version of mapNH developed in our lab (Rousselle et al. 2019) which categorizes mapped substitutions into GC-conservative (GC->GC or AT->AT) and GC-modifying (AT->GC or GC->AT) substitutions, and uses only the former to compute dN/dS ratios. Ratios obtained this way show more sampling variance, as they are obtained from smaller substitution counts. This translates in higher genomic dN/dS, as this parameter is usually close to its zero bound in exons. These rates are however supposedly impervious to gBGC.

### Controlling for sampling bias

Four Hymenoptera (Apis mellifera and the three ants), which represent nearly half the eusocial species considered, are species with published genomes. This translates into a better power for gene prediction and thus, into an over-representation of these species in the dataset. Imprecisions in dN/dS ratios estimations are in turn known to yield higher values, because the real value of this ratio in functional sequences is often close to its zero boundary. We thus applied an additional sub-sampling procedure, designed to correct for any potential bias in our estimations that could stem from variation in the quantity of information available for each species. We applied every analysis mentioned before to a reduced but complete dataset containing data only for the most represented half of the species (88 species), and only alignments containing information for each of these species (135 alignments).

### Linear modelling of dN/dS ratios

Estimated rates, corrected rates and rates obtained from the 88-species subsampled dataset were then modelled through simple linear models using the R software environment, using adult size, social status (eusocial or solitary) and membership to Anthophila as covariables. We also used this statistical setting to evaluate the effect of branch length. Short branches are known to bias dN/dS estimations upward because they yield more inaccurate and thus generally higher estimations of this parameter. The phylogenetic setting was taken into account by using phylogenetic independent contrast (Felsenstein 1981) for each variable. This was done using the pic() function in the R package ape. To try and further uncover the potential links between dN/dS ratios and life-history within Hymenoptera, we also attempted to correlate dN/dS ratios with major descriptors of parasitic type within parasitoid Hymenoptera. These descriptors were gathered from databases designed to describe the reproductive strategy of parasitoids (Traynor & Mayhew 2005a; Traynor & Mayhew 2005b, Jervis & Ferns 2011; Mayhew 2016) and are summarized in table S1. We conducted the analysis at the genus level using genus-averaged dN/dS ratios and descriptors. This was necessary because the species-level concordance between databases was too low (only 6 species in common between the genomic database and the parasitoid life-history database). We used Pearson’s linear correlation coefficient for continuous descriptors and Kruskal-Wallis tests for discrete descriptors.

Finally, we tested the correlation of dN/dS ratios with four proxies of species range. For each species (and for all known synonyms) in the sample, we queried all available occurrence points from the GBIF database, using the R package rgbif. Occurrence data was then used to calculate for classical proxies of species range. The mean latitude was calculated as a simple unweighted mean between occurrences. The maximum distance between two occurrences was calculated taking all occurrences into account, even when the species occurred on more than one continent. The circular area around occurrence was calculated by casting 100km-radius circles around each occurrence, and estimating the total land surface contained in at least one circle. The convex hull area around occurrence was calculated by estimating the total land surface contained in the smallest convex hull containing all occurrences. When a species occurred on more than one continent, a separate convex hull was used per continent.

### RELAX analyses

We used the RELAX procedure (Wertheim et al. 2014) from the HyPhy program suite (Pond, Frost, and Muse 2005) to test for the presence of a systematic relaxation of selection on branches belonging to eusocial groups (thereafter called “eusocial branches”), that is all branches descending from the ancestral node of one of the eusocial clade present in the dataset. Hyphy allows, for a specific sequence alignment, to model the distribution of dN/dS ratios along the branches of a tree. The RELAX procedure consists first in defining focal and background branches, associated with one focal and one background distribution of dN/dS ratios. It then consists in comparing a model where the two mentioned distribution are identical (null model, no differences between branch sets) to a model where the focal distribution is a power transform of the background distribution (ωf=ωbk). Relaxation of selection is inferred when the second model appears superior based on a log-ratio test (differences between branch sets), and when the focal distribution is narrower than the background distribution (k parameter estimated to be less than 1). Indeed, strong selection is thought to produce both low (close to 0) and high (greater than 1) dN/dS ratios, while neutrality should produce rates close to 1. This test thus correctly takes into account the fundamental two-sided nature of dN/dS ratios. 20 (0.61%) of the original alignments did not contain enough data to allow models necessary to the HyPhy RELAX procedure to be fitted with eusocial branches as background branches, and 17 (0.52%) of the original alignments didn’t allow the procedure with Anthophila branches as background branches.

## Supplementary

**Figure S1.**
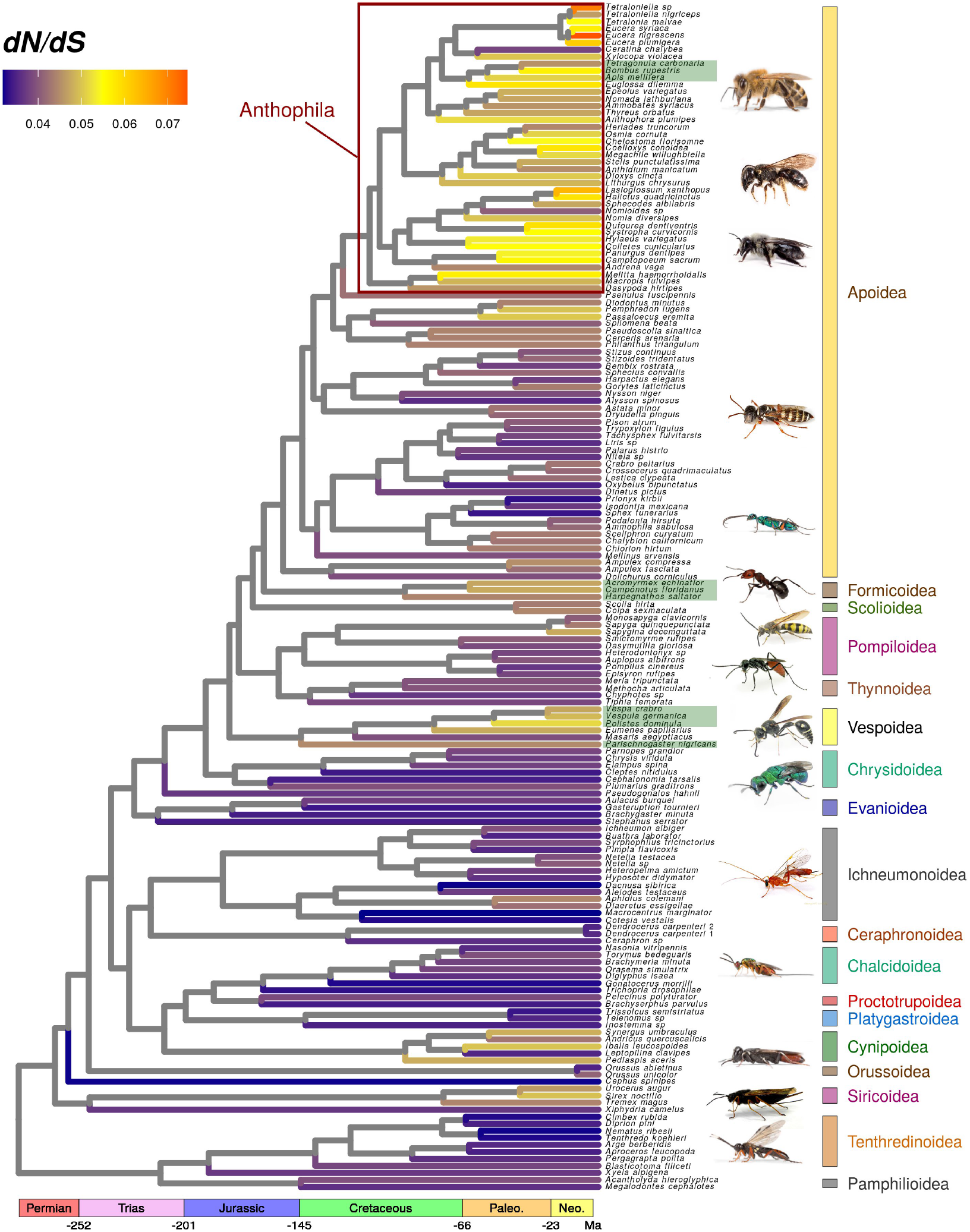
Uncorrected genomic dN/dS ratios for 169 species of Hymenoptera. dN/dS ratios estimated on terminal branches using 3241 genes and GC conservative substitutions are represented on the chronogram inferred by Peters et al. (2017). Green rectangles around labels indicate eusocial taxa.

**Figure S2.**
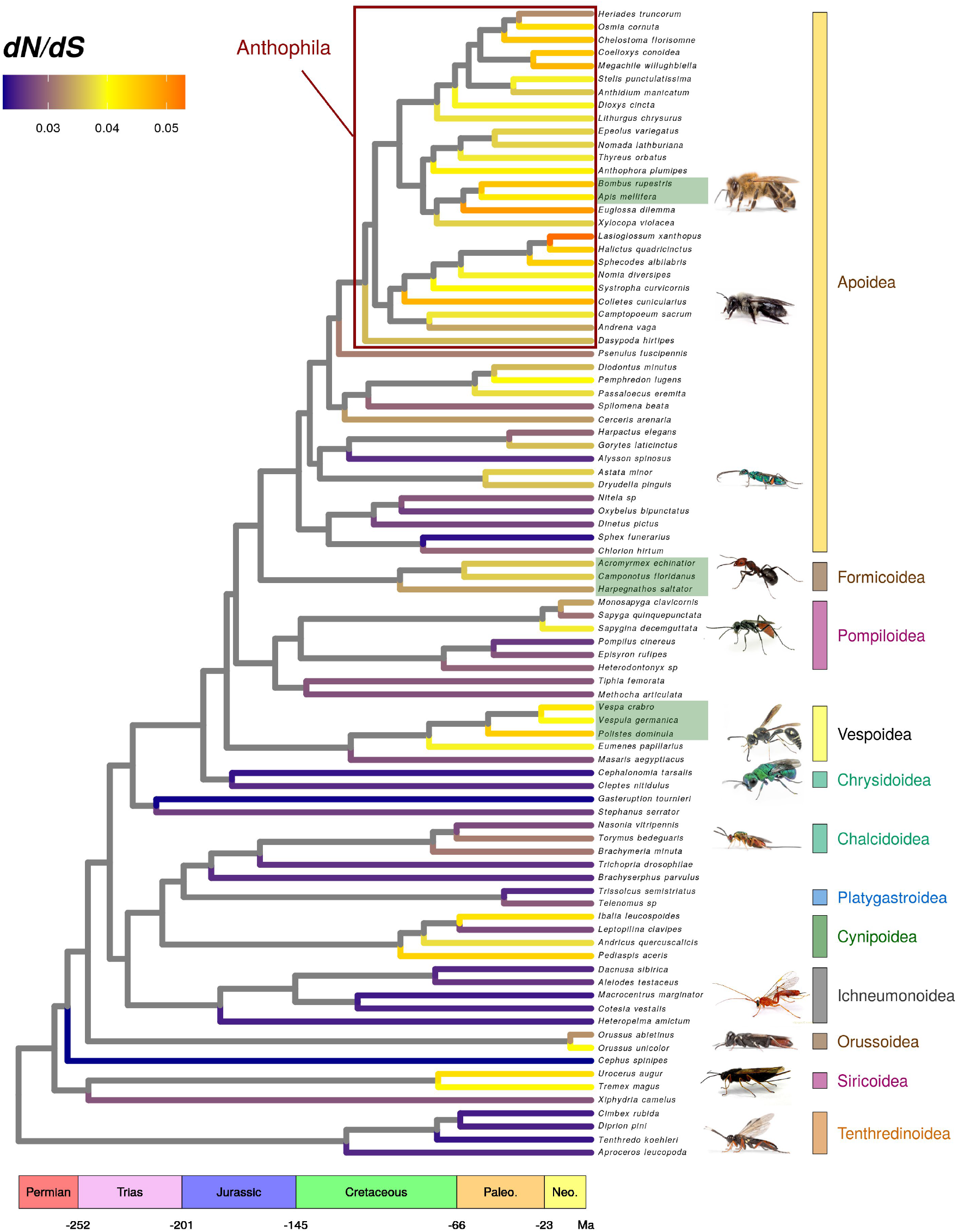
Genomic dN/dS ratios for 88 species of Hymenoptera. dN/dS ratios estimated on terminal branches using 134 genes with data for each of the displayed species are represented on the chronogram inferred by Peters et al. (2017). Green rectangles around labels indicate eusocial taxa.

**Figure S3.**
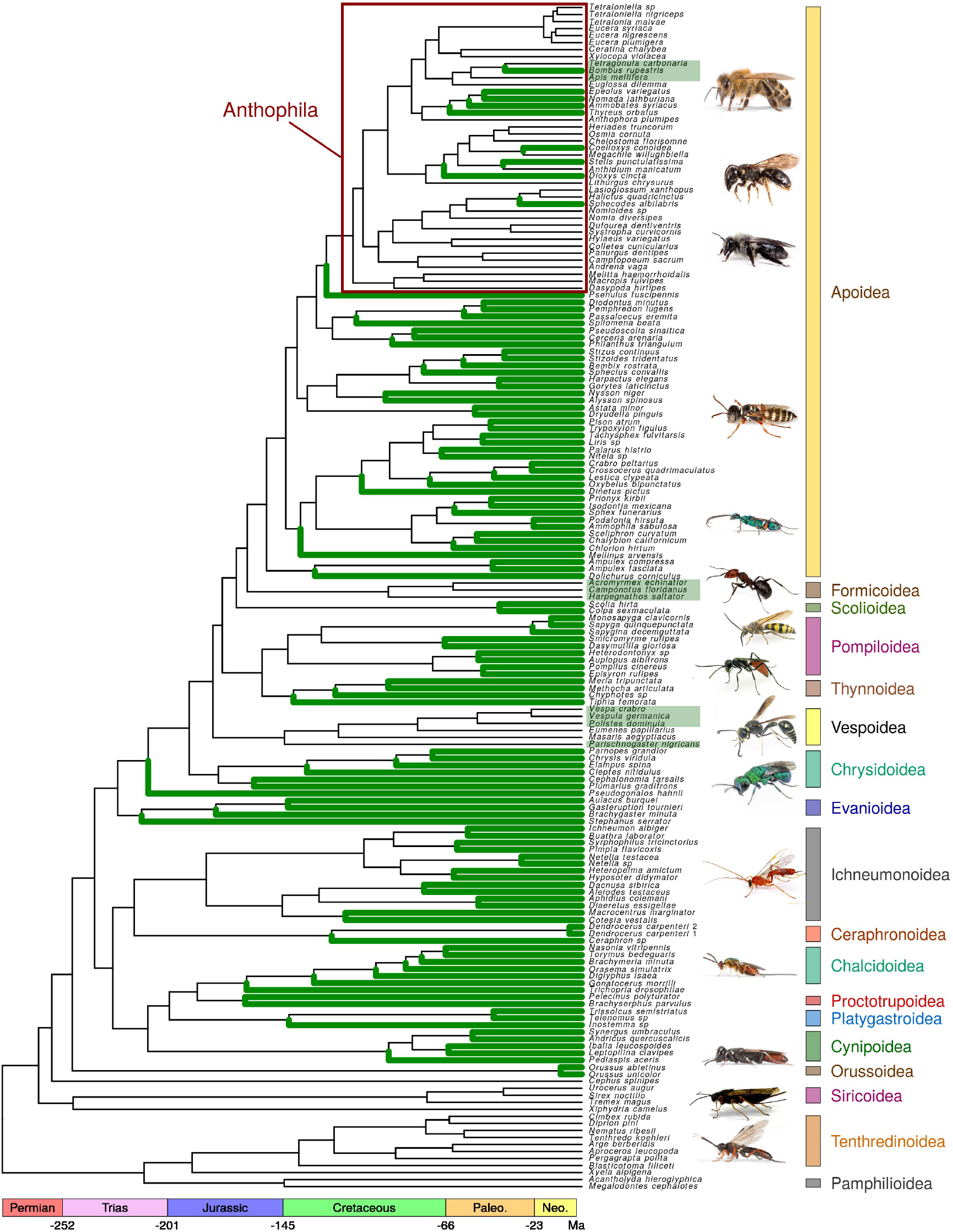
Parasitic species in the dataset. Parasitics species (parasitoids, parasites of plants and social parasites) are indicated by green terminal branches. Green rectangles around labels indicate eusocial taxa.

**Table S1.**
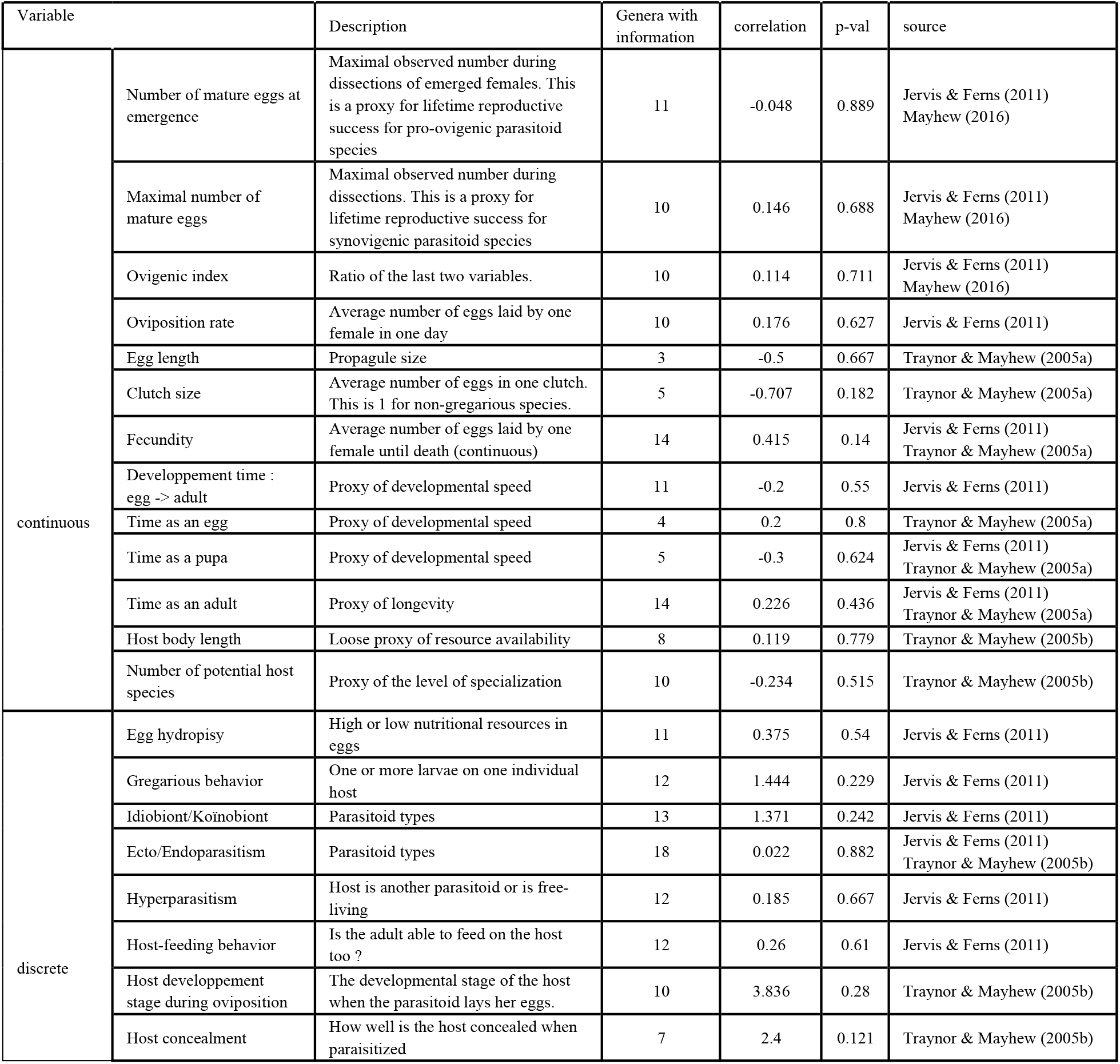
Life-history and specialisation descriptors for parasitoids. Tested variables and their description are displayed along with the value of the statistic obtained for each correlation test with corrected dN/dS ratios. Correlation tests are Spearman tests for continuous variables and Kruskal-Wallis tests for discrete variables.

**Table S2.**
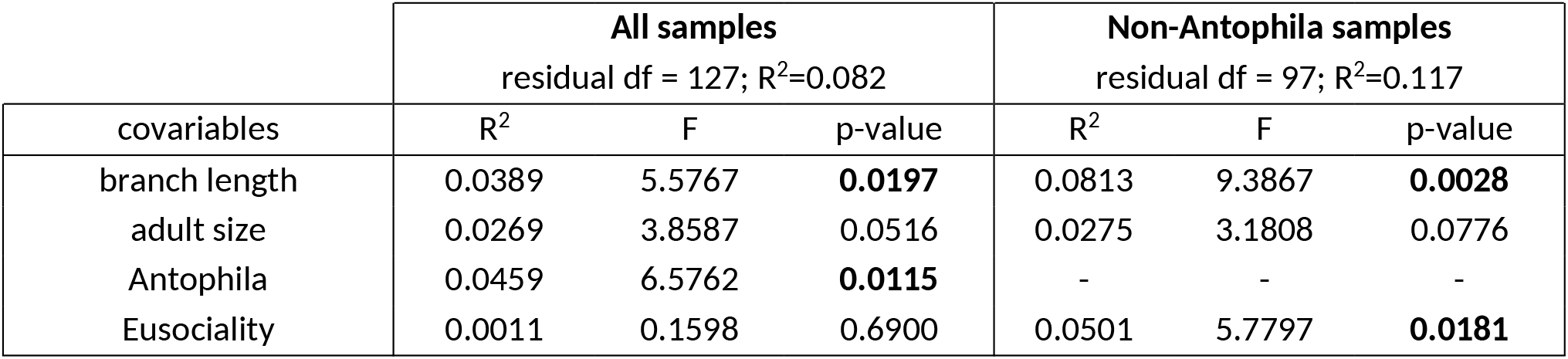
Linear modelling of uncorrected dN/dS ratios. Displayed results are obtained when simultaneously using all covariables inside a multiple linear model. Phylogenetic independent contrasts are used for all variables so as to account for phylogenetic autocorrelation.

|  | All samples<br>residual df = 127; R<sup>2</sup>=0.082 |  |  | Non-Antophila samples<br>residual df = 97; R<sup>2</sup>=0.117 |  |  |
| --- | --- | --- | --- | --- | --- | --- |
| covariables | R<sup>2</sup> | F | p-value | R<sup>2</sup> | F | p-value |
| branch length | 0.0389 | 5.5767 | **0.0197** | 0.0813 | 9.3867 | **0.0028** |
| adult size | 0.0269 | 3.8587 | 0.0516 | 0.0275 | 3.1808 | 0.0776 |
| Antophila | 0.0459 | 6.5762 | **0.0115** | - | - | - |
| Eusociality | 0.0011 | 0.1598 | 0.6900 | 0.0501 | 5.7797 | **0.0181** |

**Table S3.** Linear modelling of uncorrected dN/dS ratios in the 88-species subsampled dataset. Corrected dN/dS are obtained using GC-conservative substitutions only. Displayed results are obtained when simultaneously using all covariables inside a multiple linear model. Phylogenetic independent contrasts where used for all variables so as to account for phylogenetic autocorrelation.

|  | All samples<br>residual df = 127; R<sup>2</sup>=0.082 |  |  | Non-Antophila samples<br>residual df = 97; R<sup>2</sup>=0.117 |  |  |
| --- | --- | --- | --- | --- | --- | --- |
| covariables | R<sup>2</sup> | F | p-value | R<sup>2</sup> | F | p-value |
| branch length | 0.0281 | 2.2860 | 0.1351 | 0.0677 | 3.9381 | 0.0530 |
| adult size | 0.0283 | 2.2963 | 0.1342 | 0.0524 | 3.0525 | 0.0871 |
| Antophila | 0.0691 | 5.6131 | **0.0206** | - | - | - |
| Eusociality | 0.0237 | 1.9294 | 0.1692 | 0.0714 | 4.1567 | **0.0471** |

**Table S4.** Go terms enriched in genes supporting an intensification of selection in eusocial Hymenoptera. P-values are those of a Fisher hypergeometric test used for significance in the GO enrichment analysis, as implemented in the R package topGO (Rahnenfuhrer and Alexa 2019)

| domain | GO ID | Term | p-val |
| --- | --- | --- | --- |
| biological process | GO:0043623 | cellular protein complex assembly | 0.00011 |
|  | GO:0016043 | cellular component organization | 0.00011 |
|  | GO:0043604 | amide biosynthetic process | 0.00012 |
| molecular function | GO:0003723 | RNA binding | 0.00018 |
|  | GO:0008092 | cytoskeletal protein binding | 0.00030 |
|  | GO:0003735 | structural constituent of ribosome | 0.00083 |
|  | GO:0005488 | binding | 0.00109 |
|  | GO:0051020 | GTPase binding | 0.00113 |
|  | GO:0005085 | guanyl-nucleotide exchange factor activity | 0.00350 |
|  | GO:0017069 | snRNA binding | 0.00376 |
|  | GO:0019899 | enzyme binding | 0.00491 |
|  | GO:0005198 | structural molecule activity | 0.00500 |
|  | GO:0030246 | carbohydrate binding | 0.01089 |
|  | GO:0019904 | protein domain specific binding | 0.01286 |
|  | GO:0008536 | Ran GTPase binding | 0.02269 |
|  | GO:0003924 | GTPase activity | 0.02838 |
|  | GO:0017016 | Ras GTPase binding | 0.03243 |
|  | GO:0031267 | small GTPase binding | 0.03243 |

